# In Vivo MRI of Endogenous Remyelination in a Nonhuman Primate Model of Multiple Sclerosis

**DOI:** 10.1101/2021.10.27.466044

**Authors:** Nathanael J. Lee, Pascal Sati, Seung-Kwon Ha, Nicholas J. Luciano, Snehashis Roy, Benjamin Ineichen, Emily C. Leibovitch, Cecil C. Yen, Dzung L. Pham, Afonso C. Silva, Mac Johnson, Steven Jacobson, Daniel S. Reich

**Affiliations:** Translational Neuroradiology Section, National Institutes of Health, Bethesda, MD, USA; Viral Immunology Section, National Institutes of Health, Bethesda, MD, USA; Cerebral Microcirculation Section, Laboratory of Functional and Molecular Imaging, National Institute of Neurological Disorders and Stroke, National Institutes of Health, Bethesda, MD, USA; Department of Medicine, Massachusetts General Hospital / Harvard Medical School, Boston, MA USA; Neuroimaging Program, Department of Neurology, Cedars Sinai, Los Angeles, CA, USA; Department of Neurobiology, University of Pittsburgh, Pittsburgh, PA, USA; Section on Neural Function, National Institute of Mental Health, National Institutes of Health, Bethesda, MD, USA; Department of Neuroradiology, Clinical Neuroscience Center, University Hospital Zurich, University of Zurich, Switzerland; Center for Neuroscience and Regenerative Medicine, The Henry M. Jackson Foundation for the Advancement of Military Medicine, Bethesda, MD, USA; Vertex Pharmaceuticals Incorporated, Boston, MA USA

## Abstract

Remyelination is crucial for recovery from inflammatory demyelination in multiple sclerosis (MS). Investigating remyelination *in vivo* using magnetic resonance imaging (MRI) is difficult in MS, where collecting serial short-interval scans is challenging. Using experimental autoimmune encephalomyelitis (EAE) in common marmosets, a model of MS that recapitulates focal cerebral MS lesions, we investigated whether remyelination can be detected and characterized noninvasively. In 6 animals followed with multisequence 7-tesla MRI, 36 focal lesions, classified as demyelinated or remyelinated based on signal intensity on proton density-weighted images, were subsequently assessed with histopathology. Remyelination occurred in 5 of 6 marmosets and 51% of lesions. Radiological-pathological comparison showed high sensitivity (88%) and specificity (90%) for detecting remyelination by *in vivo* MRI. This study demonstrates the prevalence of spontaneous remyelination in marmoset EAE and the ability of *in vivo* MRI to detect it, with implications for preclinical testing of pro-remyelinating agents and translation to clinical practice.

## INTRODUCTION

Multiple sclerosis (MS) is a debilitating inflammatory demyelinating disorder affecting millions worldwide (1). MS causes dynamic changes to myelin in the central nervous system (CNS), including the quintessential focal inflammatory destruction of myelin, as well as the phenomenon of remyelination that can follow the demyelination (2-5). Although remyelination is a crucial aspect of tissue repair, and as such represents an important therapeutic target (6), most knowledge about remyelination in MS derives from postmortem studies using histochemical and electron microscopy studies. This is because investigating remyelination *in vivo* in real time is limited by imperfect discrimination on neuroimaging modalities such as magnetic resonance imaging (MRI). Furthermore, in human beings, where collecting serial short-interval scans is highly challenging, it is difficult to detect track the dynamic occurrence of remyelination. Therefore, to investigate the pathobiology of remyelination in the context of focal inflammatory demyelination, a reliable preclinical model is needed to develop techniques that can then be applied clinically.

Rodent models have been widely used to investigate various aspects of the pathobiology of demyelination. However, while toxin models in mice, including the lysolecithin and cuprizone models, display demyelination and even remyelination, they do not require involvement of adaptive immune cells (7, 8). Conversely, rodent experimental autoimmune encephalomyelitis (EAE) models, while mediated by an immune response, are often neither focal nor profoundly demyelinating. There is no known rodent model that is characterized by multifocal inflammatory demyelination in the brain that is disseminated in both space and time.

EAE in the common marmoset (*Callithrix jacchus*) is a well-recognized translational model that serves as a bridge between the rodent EAE and human MS (9-11). Not only does EAE recapitulate MS at the lesion level both radiologically and pathologically (12, 13), but lesions spontaneously remyelinate (14), as occurs in MS.

Prior studies demonstrated that certain signal changes in MRI, such as magnetization transfer ratio (MTR), correlate with remyelination in MS lesions (15-19). This has also been investigated in animal models, albeit mainly in the rodent model and in the cortex (20, 21) rather than white matter. It has also been demonstrated that partial remyelination can occur and can be localized either to specific parts of the lesion (most commonly the lesion edge) or to the whole lesion (22, 23).

Here, we studied focal white matter lesions in marmoset EAE. We utilized serial *in vivo* MRI, mainly involving proton density-weighted (PDw) and MTR sequences, to age and characterize lesions. We further analyzed the lesions using histopathology, focusing on myelin lipids and proteins, to compare and study the reliability of using various *in vivo* MRI sequences in predicting remyelination and their histopathological correlates.

## METHODS

### Marmoset EAE induction

6 marmosets (3 pairs of twins; 4 males and 2 females, ages 2–6 at baseline) were included in the study (Table 1). 2 marmosets (M#1-2) first received 100-mg of human white matter homogenate followed by an additional 200 mg after no lesions were detected by *in vivo* MRI 2 months after the initial injection (14, 24). 4 additional marmosets (M#3-6) received 200 mg of white matter homogenate, specifically from the temporal lobe, collected from autopsy. All white matter homogenates were mixed with complete Freund’s adjuvant (Difco Laboratories). Prior marmoset EAE studies have merged data from different induction protocols in an effort to make the most of this rare animal resource, and there is no evidence that lesion-level pathology differs between the two protocols used here, or between the sexes (25). In each twin pair, the first animal to develop a lesion, detected by *in vivo* MRI, received intravenous methylprednisolone (18 mg/kg/day for 5 consecutive days) with the goal to reduce the severity of inflammation, potentially allowing longer-term evaluation of the lesions; this regimen is based on treatment of acute MS relapses (26, 27). Experiments were terminated when animals became either paraplegic or lost more than 20% of their baseline body weight. For psychosocial wellbeing, all marmosets were housed with another marmoset (usually a twin counterpart). Animals were weighed and monitored on a daily basis to ensure adequate nutritional intake and physical wellbeing. All protocols were approved by the National Institutes of Neurological Disorders and Stroke Animal Care and Use Committee.

**Table 1.** Demographics and experimental information for the 6 marmosets included in this study. Immunizations used human white matter homogenate. Experiment duration corresponds to the time between immunization and terminal MRI. *, ^┼, ╫^ denote the 3 different pairs of twin animals.

| Animal | Sex | Age<br>(years) | 1st immun-<br>ization | 2<sup>nd</sup> immun-<br>ization | Corticosteroid<br>treatment | Experiment<br>duration<br>(weeks) |
| --- | --- | --- | --- | --- | --- | --- |
| M#1* | Male | 3.7 | 100 mg | 200 mg | Yes | 40 |
| M#2* | Male | 3.7 | 100 mg | 200 mg | No | 54 |
| M#3<sup>†</sup> | Male | 6.2 | 200 mg | - | Yes | 61 |
| M#4<sup>†</sup> | Male | 6.2 | 200 mg | - | No | 60 |
| M#5<sup>‡</sup> | Female | 2.8 | 200 mg | - | Yes | 18 |
| M#6<sup>‡</sup> | Female | 2.8 | 200 mg | - | No | 16 |

### Marmoset *in vivo* MRI

All 6 marmosets were scanned on a weekly basis under anesthesia, as previously described (12, 14, 24). We used PDw contrast, which is sensitive to demyelination (28), to visualize lesions *in vivo*. T1-weighted (T1w), T2-weighted (T2w), T2*-weighted (T2*w) and MTR contrasts were also obtained (Figure 1). T1w contrast was repeated after injection of gadolinium-based contrast agent (gadobutrol, 0.3 mg/kg) to visualize enhancing lesions. Specific sequence parameters for the different MRI contrasts are listed on Table 2. Images were post-processed using an in-house pipeline, including intensity correction, image cropping, multi-contrast registration, skullstripping, and intensity normalization (Supplementary Figure 1). To minimize harm and pain in animals, all procedures, including intravenous access for gadolinium injection, were done under anesthesia. For post-anesthesia recovery, animals were gently woken up using warm blankets, and returned to their respective housing only after the animals were back to pre-anesthesia baseline, including spontaneous breathing, physical activities, and intractability.

**Figure 1.**
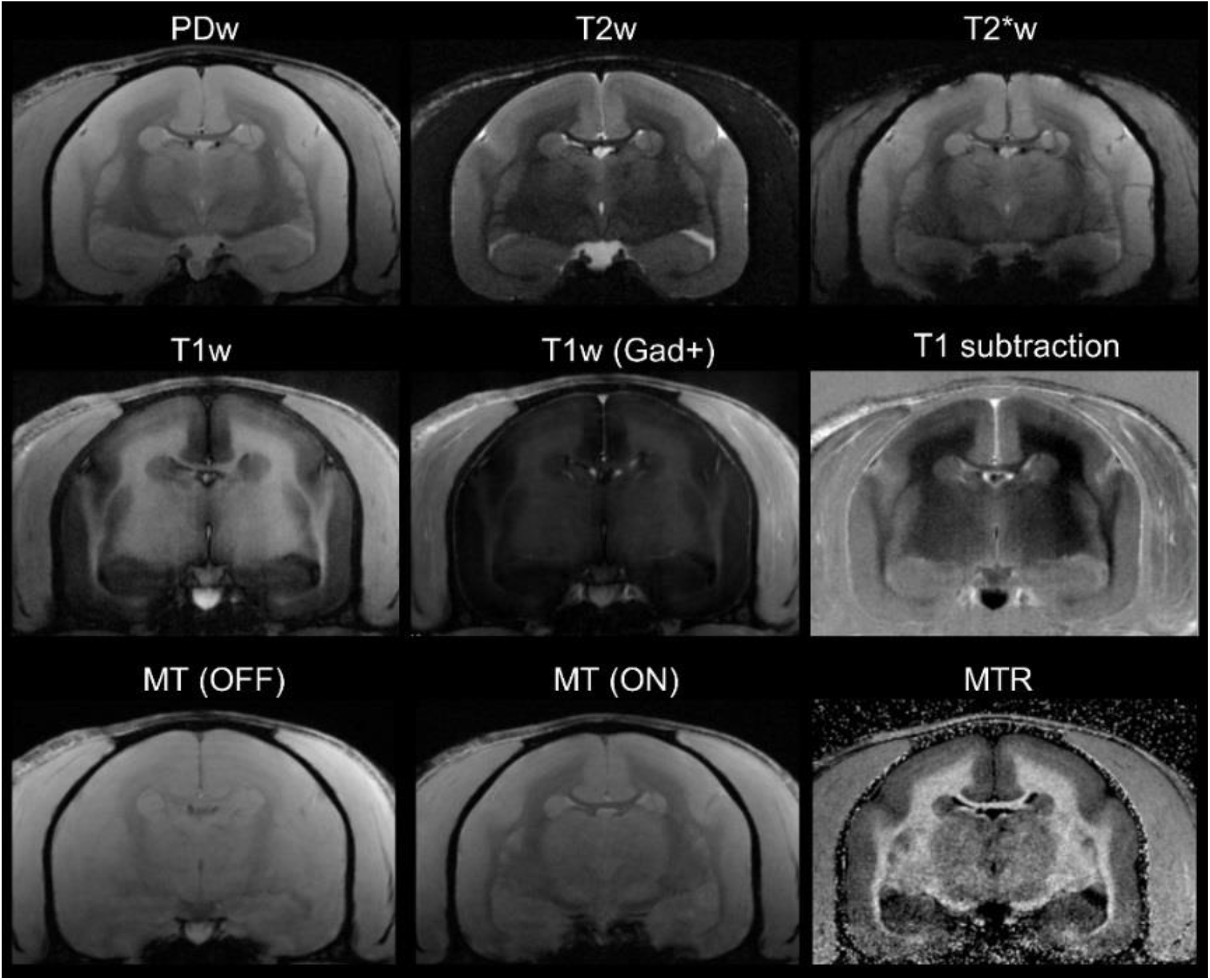
*In vivo* (baseline) multicontrast marmoset brain MRI. T1 subtraction image was obtained from subtracting pre-gadolinium T1w from post-gadolinium T1w images. MTR image derived voxelwise as *(M*_*0*_ *– M*_*SAT*_*)/ M*_*0*_. Animal M#5.

**Table 2.** Main parameters used for the different MRI contrasts acquired *in vivo*. FOV = field of view; TR = repetition tine; TE = echo time; TI = inversion time; ETL = echo train length (or RARE factor); Exc. Pul. = excitation pulse; Ref. Pul. **=** refocusing pulse; Pre. Pul. = preparation pulse, FA = flip angle; NEX = number of repetitions; AT = acquisition time; RARE = rapid acquisition with relaxation and enhancement; MDEFT = modified drive equilibrium Fourier transform; MGE = multi gradient echo. ^┼^sequence was performed twice: with (M_SAT_) and without (M_0_) the MT pre-pulse.

| **MRI contrast** | **PDw** | **T1w** | **T2w** | **T2\*w** | **MTR<sup>†</sup>** |
| --- | --- | --- | --- | --- | --- |
| Sequence | 2D RARE | 2D MDEFT | 2D RARE | 2D MGE | 3D MGE |
| FOV (mm) | 32 x 24 | 32 x 24 | 32 x 24 | 32 x 24 | 38.4 x 38.4 x 36 |
| Matrix | 214 x 160 | 214 x 160 | 214 x 160 | 214 x 160 | 256 x 256 x 36 |
| Number of slices | 36 | 36 | 36 | 36 | 36 |
| Slice thickness (mm) | 1 | 1 | 1 | 1 | 1 |
| TR (ms) | 2300 | 12.5 | 8000 | 2150 | 20 |
| TE (ms) | 16 | 4.2 | 72 | 18 | 5 |
| TI (ms) | N/A | 1200 | N/A | N/A | N/A |
| ETL | 1 | N/A | 4 | N/A | N/A |
| Exc. Pul. (shape, FA) | Sinc3, 90° | Sinc3, 12° | Sinc3, 90° | Sinc3, 70° | Sinc3, 10° |
| Ref. Pul. (shape, FA) | Sinc3, 180° | N/A | Sinc3, 180° | N/A | N/A |
| Pre. Pul. (type, shape, FA, offset) | N/A | Exc./Inv., Sech, 90°/180° | N/A | N/A | MT, Gauss, 1500 Hz |
| NEX | 1 | 1 | 2 | 1 | 2 |
| AT | 7 min 40 sec | 6 min 56 sec | 13 min 20 sec | 7 min 10 sec | 7 min 58 sec |

### Histopathology of EAE lesions

Marmoset brains were collected immediately after death once the animals met the study end-point. Brain tissue was processed using formalin-fixation and paraffin-embedding and subsequent histopathological staining, as described previously (14, 24). Postmortem histological processing failed for animal M#1’s brain. For visualizing myelin, Luxol fast blue (LFB) staining with periodic acid Schiff (PAS) counterstain and immunohistochemistry for myelin proteolipid protein (PLP) were used. For characterizing inflammation, edema, gliotic activity, hematoxylin and eosin (H&E) and immunohistochemistry for ionized calcium-binding adaptor molecule (Iba1) were used. Oligodendrocytes and oligodendrocyte precursor cells (OPC) were assessed with aspartoacylase (ASPA) and oligodendrocyte transcription factor (Olig2) double staining (mature oligodendrocytes were ASPA and Olig2 positive; OPCs were Olig2 positive but ASPA negative). For axon staining, Bielschowsky’s silver method was used. Briefly, deparaffinized slides were covered with 20% AgNO_3_ and incubated at 40 °C inside a dark chamber for 30 minutes. Slides were washed and placed in ammonia silver solution, prepared by adding concentrated ammonium hydroxide drop-by-drop into AgNO_3_ until brown precipitate disappeared, at 40

°C for 30 minutes. Developer working solution was added to the slides, made with developer stock solution (37–40% formaldehyde, citric acid, and nitric acid), ammonium hydroxide, and distilled water. After all incubations, slides were washed with 1% ammonium hydroxide, washed in distilled water, and treated with 5% sodium thiosulfate solution. Detailed immunohistochemical information is provided in Supplementary Table 1.

### *In vivo* MRI analysis of EAE lesions

Focal marmoset EAE lesions in the white matter were detected on PDw images using an automated convolutional neural network (CNN)-based segmentation algorithm (29, 30). The network was trained using PDw MRI images from 3 EAE marmosets along with binary manual segmentation of lesions. An atlas consists of a pair of postprocessed PDw images (baseline and a time-point) and the binary lesion segmentation of that time-point. The baseline is assumed to be lesion free. Once the network was trained, it was applied to the serial PDw images collected on our 6 animals (Supplementary Figure 2), and lesion masks were automatically generated. Any lesion smaller than 4 voxels (0.0225 μl) was removed from analysis. For temporal progression computation, a lesion at timepoint *t*=*t*_1_ was identified as the same lesion at timepoint *t*=*t*_2_ if they overlapped by at least 4 voxels in 3D. All automated lesion segmentations were reviewed by an experienced rater. For subsequent MRI analysis of lesion trajectories, including intensity changes over time, all lesions were identified on every scan of each marmoset, and the average intensities were calculated using the automatically segmented lesions. For pre-lesion timepoints, a region of interest (ROI) was drawn manually and positioned in the normal appearing white matter area where the lesion later appeared.

### EAE lesion characterization, categorization, and comparison

EAE lesions were characterized independently and separately based on *in vivo* MRI characteristics and histopathological analyses (2 separate experienced raters for MRI, and 2 experienced raters for histopathological analysis, blinded to each other). The lesions were categorized as follows: acute demyelinating, chronic demyelinated, and remyelinated. Once the lesions were independently analyzed, the results were directly compared for statistical analyses as described below.

### Statistical analysis

To evaluate the sensitivity and specificity of *in vivo* MRI detection of demyelination or remyelination, relative to by histopathology, we created confusion matrices and calculated true or false positive and negative rates. For interrater reliability of using MRI to predict remyelination, Cohen’s kappa was calculated. To test the effects of corticosteroid treatment and sex on remyelination, the point biserial correlation model was used.

## RESULTS

### Lesion characterization and categorization

Typical acute demyelinating lesions were hyperintense on PDw images until the terminal scan, grew rapidly to several cubie millimeters immediately after establishment, and then showed minimal lesion volume change over time (example in Figure 2A–B). Post-gadolinium T1w and subtraction images also showed active gadolinium enhancement, as described previously (24, 31). On histopathology, LFB-PAS showed prominent demyelination, with PAS^+^ macrophages (Figure 2C). PLP staining demonstrated demyelination with myelin debris at the lesion border. Lesions harbored prominent Iba1^+^ cell infiltration, loss of axons on Bielschowsky silver staining, and loss of oligodendrocytes on ASPA/Olig2. H&E staining showed edema marked by irregular clear space.

**Figure 2.**
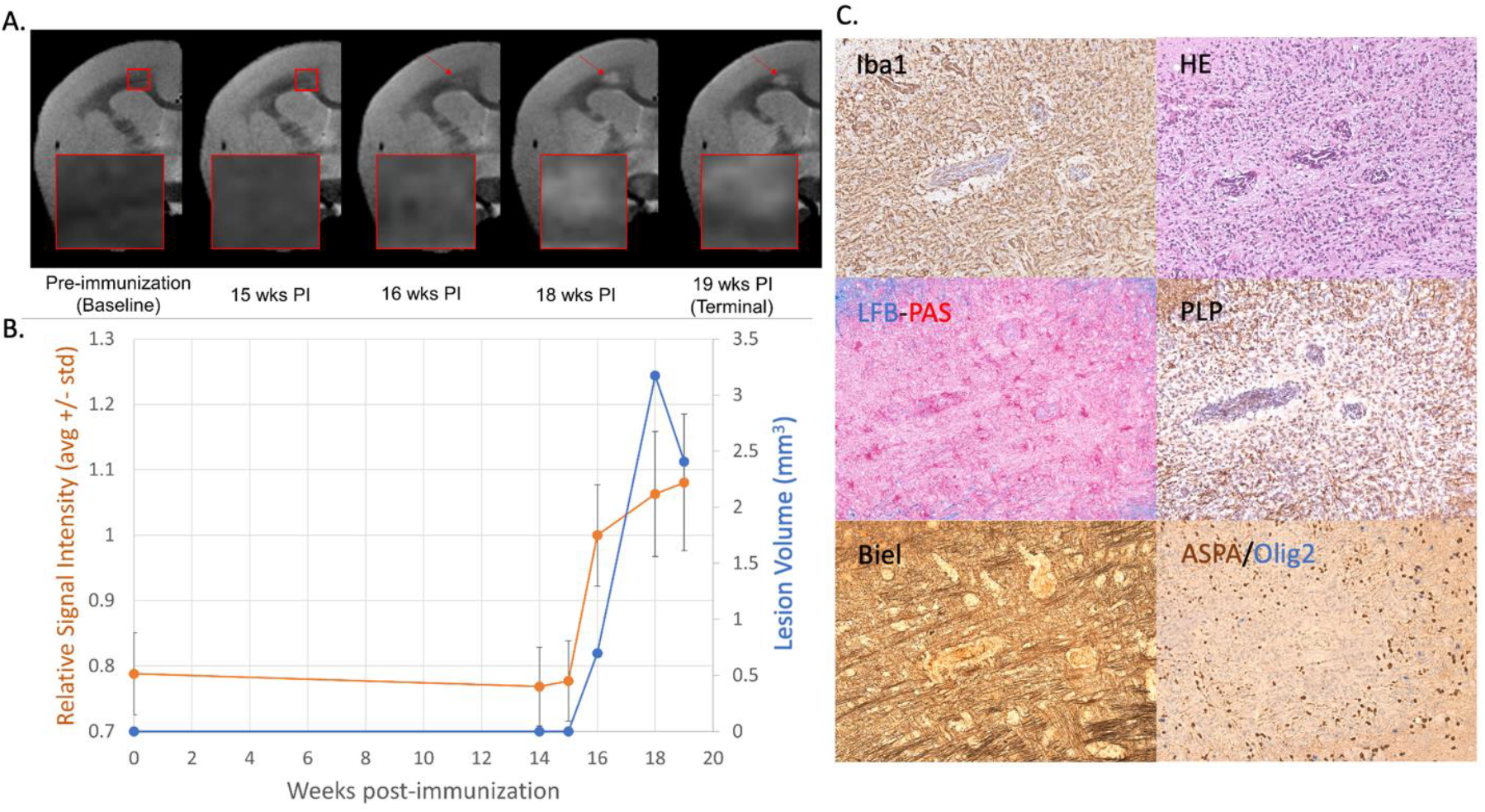
Acute demyelinating EAE lesion with persistent hyperintensity on PDw MRI and active inflammatory demyelination on histopathology. (A) *In vivo* MRI of proton density-weighted (PDw) acquired before EAE induction (baseline) and at various timepoints leading up to terminal scan. Images were processed as described in Methods. Red arrows = focal white matter lesion first detected 16 weeks after immunization, which persisted through the terminal MRI. Red boxes = magnified insets. (B) Temporal evolution of volume (blue line) and PDw signal (orange line) of the segmented lesion. Lesion signal intensity is normalized to gray matter signal intensity, which was set to 1. (C) Histochemical panel at 200X magnification of the same lesion, demonstrating active inflammatory demyelination and edema, loss of oligodendrocytes, and axon loss. Hematoxylin counterstaining used for PLP and Iba1. Lesion selected from M#6.

Typical chronic demyelinated lesions demonstrated long-lasting hyperintensity on PDw images until the terminal scan with resolution of gadolinium enhancement on T1w images (Figure 3A–B). PLP staining showed sharp lesion borders with complete demyelination on LFB-PAS (Figure 3C). Lesions contained clusters of Iba1^+^ cells. ASPA/Olig2 staining revealed loss of both mature oligodendrocytes and OPC. H&E staining showed less edema compared to acute demyelinating lesions, and Bielschowsky silver stain demonstrated loss of axons.

**Figure 3.**
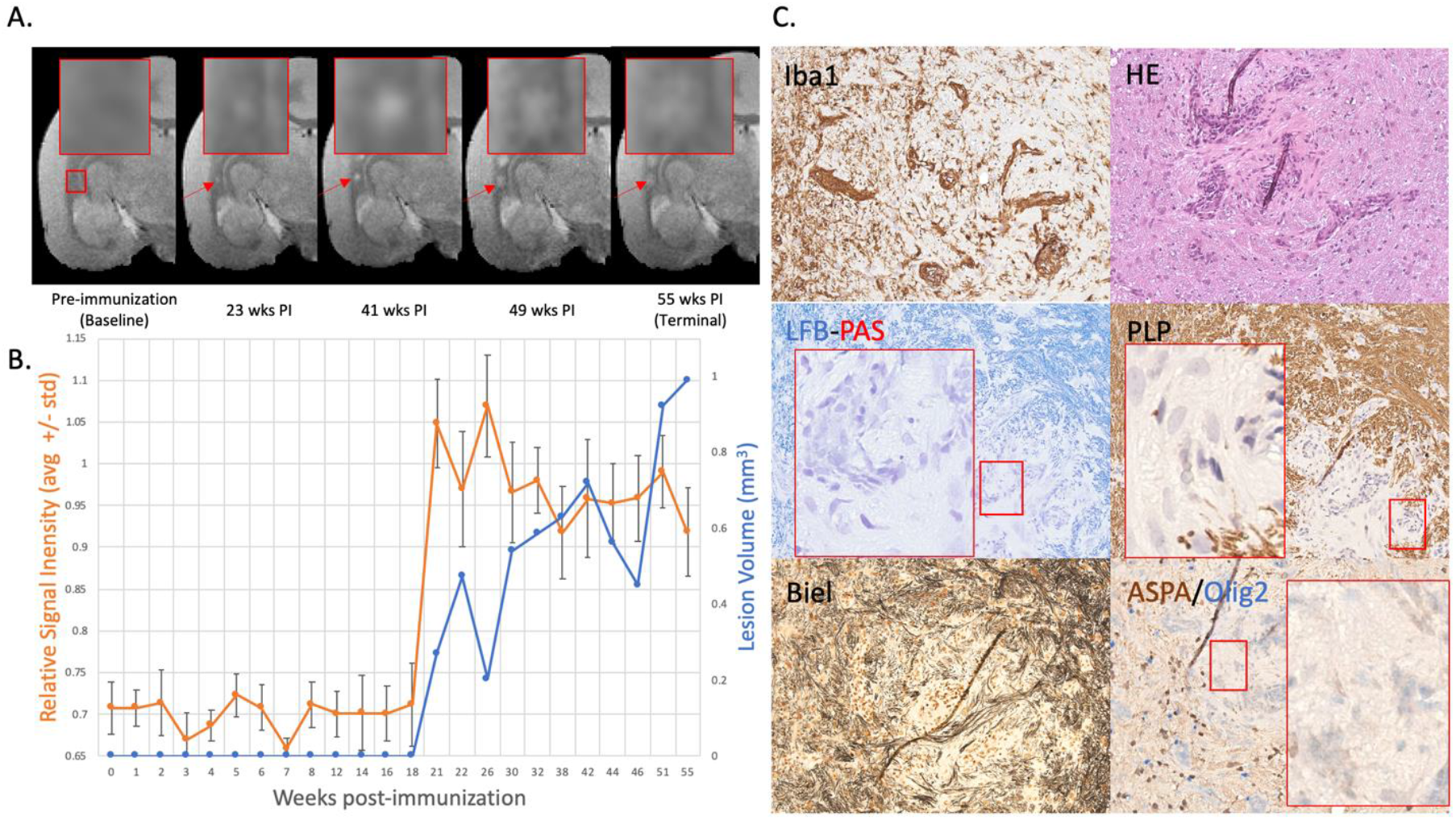
Chronic demyelinated EAE lesion with long-lasting hyperintensity on serial PDw MRI and complete loss of myelin on histopathology. (A) *In vivo* MRI of proton density-weighted (PDw) acquired before EAE induction (baseline) and at various timepoints leading up to terminal scan. Images were processed as described in Methods. Red arrows = focal white matter lesion first detected 23 weeks after immunization, which persisted until the terminal MRI. Red boxes = magnified insets. (B) Temporal evolution of volume (blue line) and PDw signal (orange line) of the segmented lesion, with acute and persistent increase in signal intensity and increasing lesion volume. Lesion signal intensity is normalized relative to gray matter signal intensity, which was set to 1. (C) Histochemical panel at 200X magnification of the same lesion showing loss of myelin, axons, and oligodendrocytes, as well as persistent inflammation. Red boxes = magnified insets. Hematoxylin counterstaining used for PLP and Iba1. Lesion selected from M#3.

Typical remyelinated lesions were initially hyperintense on PDw images, then returned to near isointensity over time (Figure 4A–B). On histopathology, most areas demonstrated nearly normal myelin structure, with occasional wave-form morphology with varying orientation (Figure 4C). Inflammatory cells were less prominent, and Bielschowsky staining showed denser normal axon structure. ASPA/Olig2 staining also demonstrated repopulation of oligodendrocytes and OPC.

**Figure 4.**
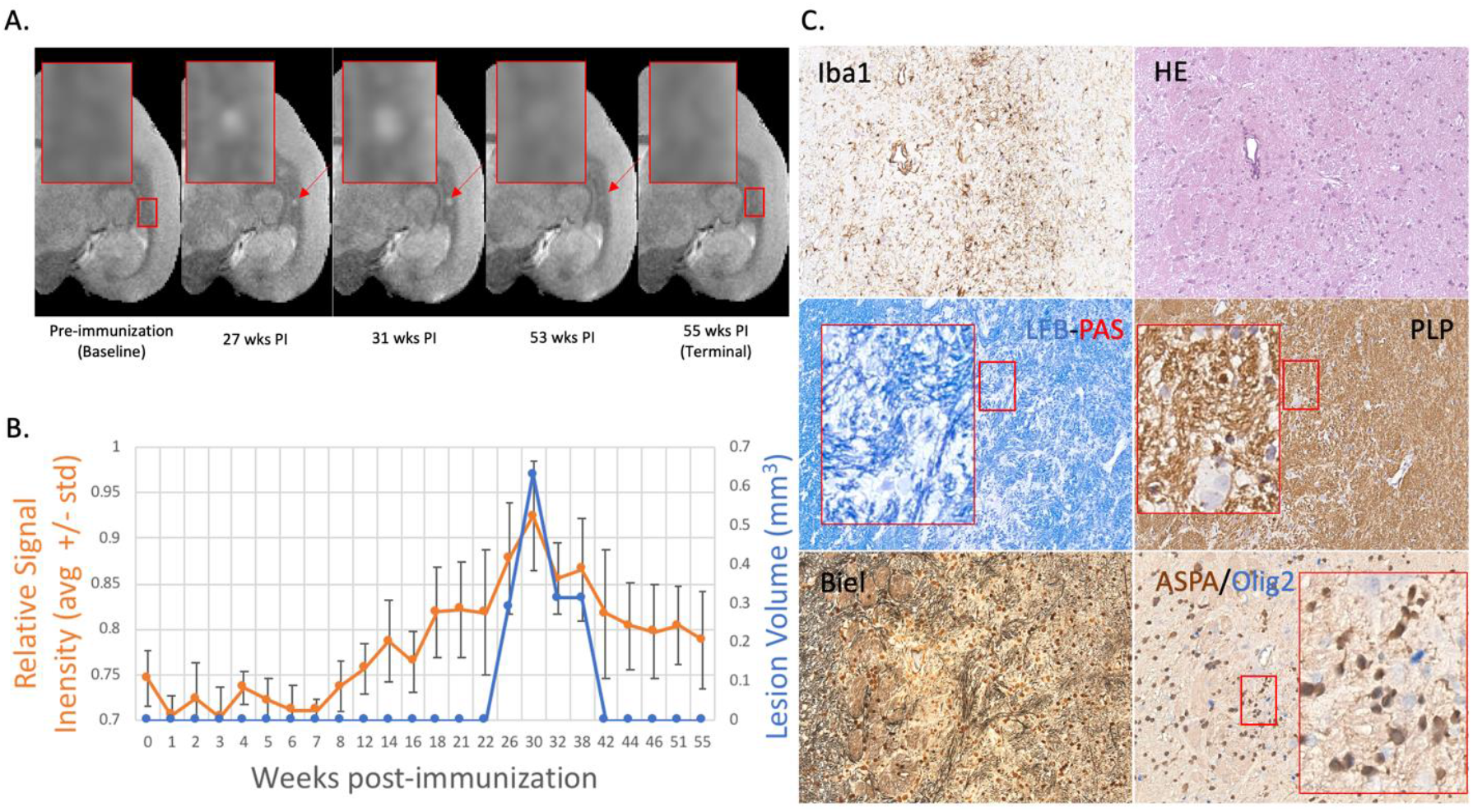
Remyelinated EAE lesion with initial hyperintensity that returned to isointensity. (A) *In vivo* proton density-weighted (PDw) MRI acquired before EAE induction (baseline) and at various timepoints leading up to the terminal scan. Images were processed as described in Methods. Red arrows = focal white matter lesion first detected 27 weeks post immunization (PI), which appeared to subsequently resolve on MRI. Red boxes = magnified insets. (B) Temporal evolution of volume (blue line) and PDw signal (orange line) of the segmented lesion. Signal intensity peaked around 31 weeks PI and returned to baseline by 42 weeks PI. Lesion signal intensity is normalized relative to gray matter signal intensity, which was set to 1. (C) Histochemical panel at 200X magnification of the same lesion showing sparse myelin and presence of both oligodendrocytes on OPC, as well as less inflammation and edema than other lesion types. Axons appeared closer to within normal limits compared to demyelinated lesions. Red boxes = magnified insets. Hematoxylin counterstaining used for PLP and Iba1. Lesion selected from M#3.

### Spontaneous remyelination is common in marmoset EAE

Using *in vivo* MRI only, 43 focal white matter lesions were detected in the 6 EAE marmosets (Table 3). Of these 43 lesions, 22 (51%) were classified as remyelinated, 7 as acute demyelinating (16%), and 14 as chronic demyelinated (33%). 4 of the 6 animals demonstrated remyelinated lesions (M#1–4), whereas the other two only had acute demyelinating and chronic demyelinated lesions (M#5–6).

**Table 3:** Classification of focal lesions by MRI.

| Animal | Acute Demyelinating | Chronic Demyelinated | Remyelinated | Total |
| --- | --- | --- | --- | --- |
| M#1 | 0 | 1 | 6 | 7 |
| M#2 | 0 | 1 | 9 | 10 |
| M#3 | 0 | 1 | 5 | 6 |
| M#4 | 0 | 1 | 2 | 3 |
| M#5 | 5 | 4 | 0 | 9 |
| M#6 | 2 | 6 | 0 | 8 |
| Total | 7 | 14 | 22 | 43 |

**Table 4:**
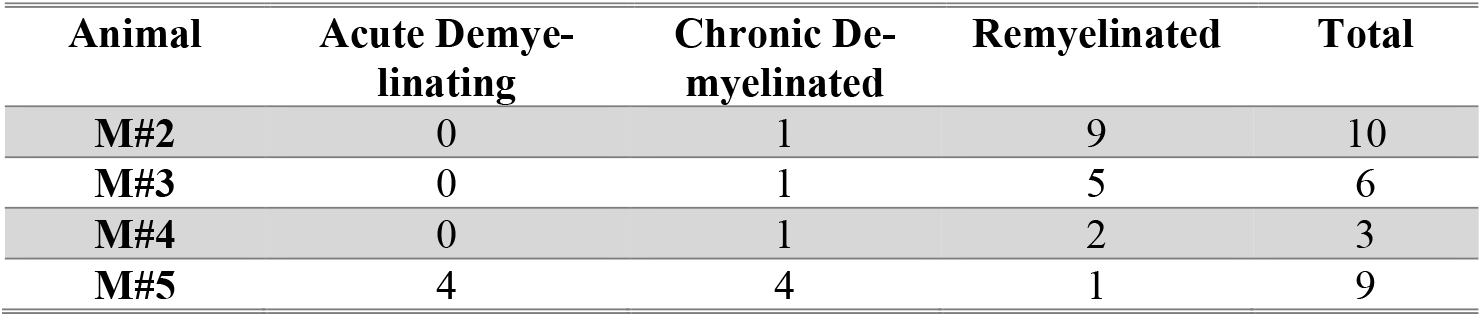
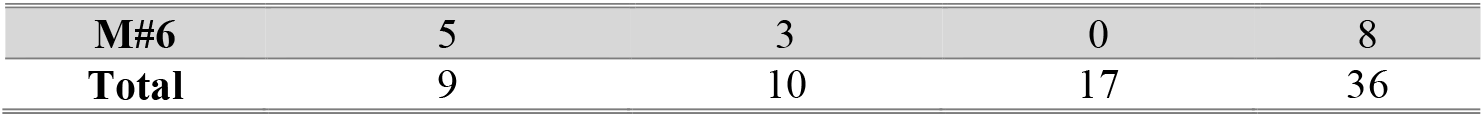
Classification of focal lesions by histopathology.

Based on the histopathological classification criteria, 36 lesions were identified in the 5 animals with postmortem tissue. 17 of the 36 lesions (47%) were classified as remyelinated, 9 as acute demyelinating (25%), and 10 as chronic demyelinated (28%). All 3 animals with putative remyelinated lesions on MRI (M#2–4) had had remyelinated lesions on histopathology, and M#5 had a single remyelinated lesion not seen on MRI; M#6 only demonstrated acute and chronic demyelinated lesions, consistent with the MRI.

### Remyelination is reliably detected by *in vivo* MRI

Of the 36 focal white matter lesions identified on both *in vivo* PDw MRI and histopathology, classification was concordant for 26 lesions (72%). Of these, 5 (19%) were acute demyelinating, 6 (23%) were chronically demyelinated, and 15 were remyelinated (58%). When the lesions were grouped by myelination status only (remyelinated vs. demyelinated), *in vivo* PDw MRI correctly predicted 32 out of 35 lesions, showing 89% agreement (Table 5). Using histopathology as the gold standard, there was 88% sensitivity and 90% specificity for detecting remyelination. Interrater reliability for use of serial PDw MRI to detect remyelination was 94%, with Cohen’s kappa of 0.89.

**Table 5:** Confusion matrix for PDw MRI classification of demyelinated vs. remyelinated lesions compared to histopathology.

| **n=36** | **Predicted demyelination by MRI** | **Predicted remyelination by MRI** |  |
| --- | --- | --- | --- |
| **Actual demyelination by histopathology** | 17 | 2 | 19 |
| **Actual remyelination by histopathology** | 2 | 15 | 17 |
|  | 19 | 17 |  |

MTR was separately used to classify lesions, in order to compare its utility to predict myelin status with PDw. The sensitivity and specificity for detecting remyelination, using histopathology as the standard, were 82% and 79%, respectively, both lower than for PDw.

### Remyelination is a time-specific event in marmoset EAE

Based on our frequent longitudinal MRI-based sampling and aging of EAE lesions and analysis of signal intensity, with comparison to histopathology, we found that inflammation and demyelination are the primary components of lesion pathophysiology in lesions <5 weeks old, corroborating our previous work (14, 24). The earliest remyelination detected was 5 weeks after initial lesion detection on *in vivo* MRI, although most remyelinated lesions were >7 weeks old. To estimate the rate of remyelination, we performed an initial estimate in 4 focal lesions followed closely by MRI, based on continuous decrease in signal intensity on PDw, and validated by histopathology (Figure 5). These data suggest that remyelination period typically takes 4–9 weeks.

**Figure 5:**
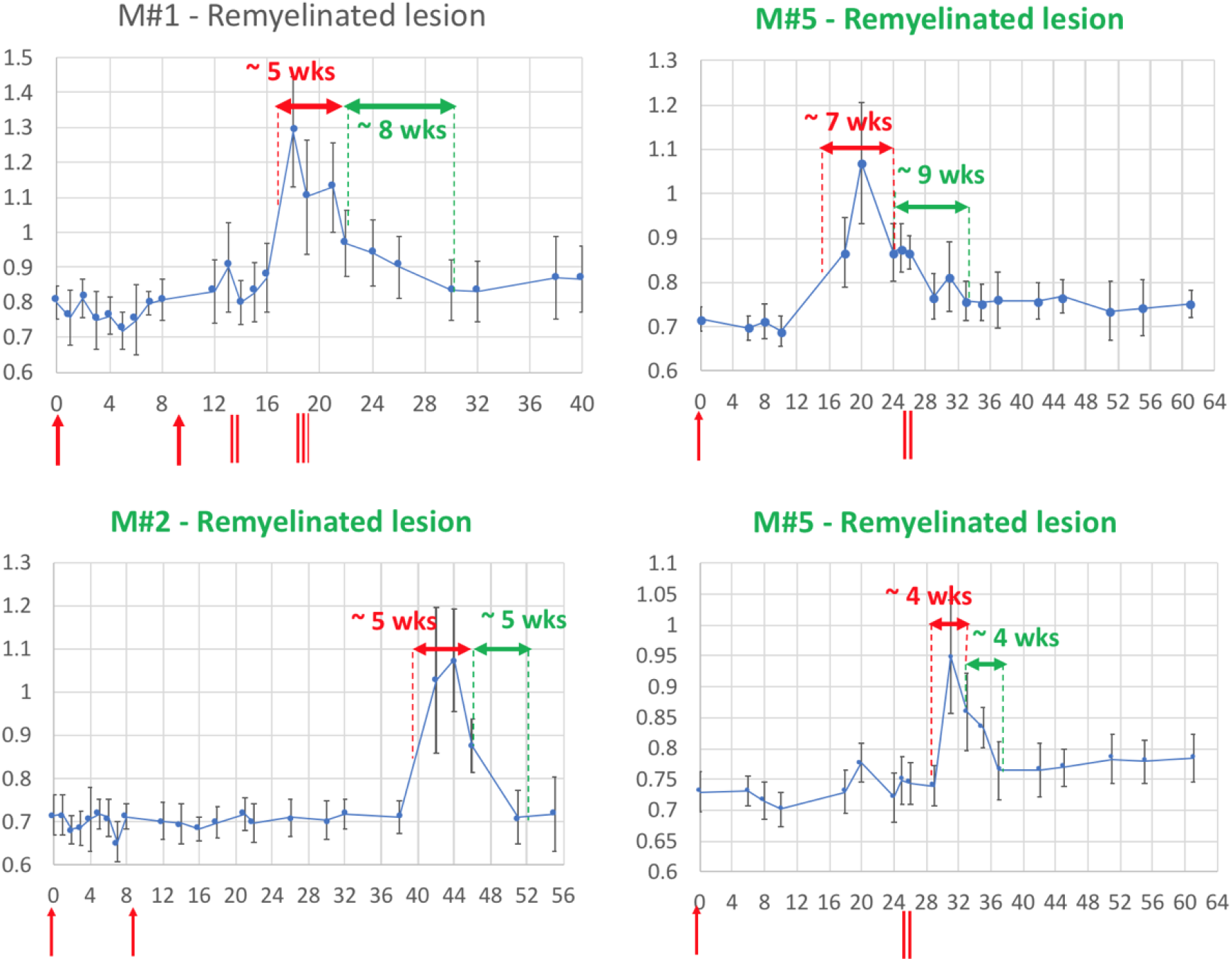
Evolution of *in vivo* PDw MRI signal intensity of remyelinating EAE lesions shows the typical 4–9-week time course of remyelination. Vertical axis: mean PDw signal intensity relative to gray matter. Horizontal axis: weeks post-immunization. Blue line corresponds to mean PDw signal (with standard deviation) of the segmented lesion as visible on MRI and quantified using a region of interest drawn manually and located in the normal appearing white matter area before the lesion appeared. Normal white matter displays an average normalized signal intensity of 0.65–0.75. Vertical red arrows indicate EAE immunization. Vertical red bars indicate corticosteroid treatment. Horizontal red double arrows indicate the estimated period of demyelination. Horizontal green double arrows indicate the estimated period of remyelination based on the downward slope of intensity measurement. Green titles indicate that the lesion subtype was confirmed with histopathology counterpart. M# corresponds to animal number in Table 1.

### Remyelination is independent of corticosteroid administration

3 of the 6 marmosets were given corticosteroids for 5 consecutive days to determine whether this treatment might change the course of the repair process. Our analysis shows that marmosets treated with steroids had no difference in the prevalence of remyelinated lesions. Based on serial PDw MRI, 11 of 22 lesions (50%) were remyelinated in corticosteroid-treated marmosets, whereas in untreated marmosets, 11 of 21 lesions (52%) were remyelinated. Based on histopathological analysis, 6 of 15 lesions (40%) in corticosteroid-treated marmosets were remyelinated, and 11 of 21 (52%) lesions in untreated marmosets were remyelinated. The point biserial correlation model analysis showed that steroid administration had no significant correlation with remyelination (*p* = 0.8). The average experiment duration also did not differ significantly (40 weeks in treated and 43 weeks in untreated marmosets).

### Remyelination may be sex-dependent

Using the point biserial correlation model analysis, there was significant correlation (r^2^ = 0.94, p = 0.013) favoring males (n = 4) over females (n = 2) for prevalence of remyelinated lesions.

## DISCUSSION

In this study, we investigated the remyelination process in focal brain lesions of marmoset EAE, a relatively faithful MS model, using high-resolution serial *in vivo* MRI and histopathology. Our main results show that lesion remyelination is common phenomenon in this model. Indeed, 5 of the 6 marmosets displayed evidence of remyelination on either MRI or histopathology, and nearly half of the 43 lesions identified on MRI showed signs of remyelination. In humans, the prevalence of remyelination is thought to be variable across MS patients and phenotypes, though it is not well estimated (4, 5, 23). Interestingly, the two animals that did not show remyelination according to *in vivo* MRI, M#5 and M#6, had the shortest disease course (16-18 weeks) compared to the other 4 animals (40-61 weeks), suggesting that these animals may have had a more aggressive course of the disease — although a single remyelinated lesion was detected histopathologically in M#5.

Our characterization of remyelinated lesions indicated diminished inflammation, repopulated myelin sheaths in different morphology compared to normal-appearing white matter, and repopulation of both mature oligodendrocytes and OPC. Remyelination in MS lesions also directly correlates with the presence of intralesional oligodendrocytes (4, 32, 33). Assessment of acute demyelinating and chronic demyelinated lesions showed fewer oligodendrocytes, and in this histopathological setting, the lesions were hyperintense on *in vivo* PDw MRI. Taken together, these data suggest that lesions that initially present as hyperintense foci on PDw MRI and subsequently return to isointensity are in fact remyelinated. In this way, signal intensity time courses can be used to infer lesion myelination status in the setting of early lesion development and recovery, even though PDw MRI itself is not pathologically specific.

The time course of marmoset EAE lesion development and repair is relatively stereotyped, as confirmed here. Previous work from our group has shown that once lesions form, the period of demyelination lasts around 6 weeks (14, 24), consistent with data from this study (4–7 weeks; see Figure 5). On the other hand, remyelination occurs over a period of 4–9 weeks after peak lesion signal intensity and volume, without much delay. This finding is in parallel with previous studies that show that remyelination is especially prominent at early stages of MS lesion development, but rarer after years of disease duration in chronic lesions (5).

Our finding that remyelination can be detected on serial *in vivo* MRI with high sensitivity and specificity, especially on PDw MRI, corroborates and expands our previous finding that PDw MRI is sensitive to myelin status changes (24, 34). Those prior studies primarily focused on demyelination but also included limited data on remyelination, with electron microscopy showing thinned myelin sheaths in remyelinated axons (14). Here, we added comparison of PDw and MTR, since MTR has been used to detect remyelination *in vivo* (15, 16, 35), finding that PDw was more sensitive and specific for identifying remyelination. Several groups have reported that MTR may be low in remyelinated lesions compared to normal-appearing white matter because of the presence of incomplete or morphologically different myelin sheaths in previously demyelinated regions (3, 35). Unlike MTR, which suffers from signal-to-noise reductions due to its calculation as a voxel-wise division of random variables, PDw signal intensity is directly measured by the MRI system, and thus PDw may prove more useful for simply discriminating the presence or absence of remyelination, and for characterizing its time course (as was done here).

To determine whether corticosteroid treatment alters the course of lesion repair, half of the marmosets (for each twin pair, the one that showed lesions first) received 5-day treatment courses. Our data show that the difference in the prevalence of remyelinated lesions, and the duration of the event, did not differ by treatment group. One possible explanation for this result is that the early initiation of corticosteroid treatment (as soon as the first lesion was detected), resulting in treatment completion before lesions were even 1 week old, might have been premature: the initial demyelination period typically lasts 4–7 weeks before remyelination ensues. It is also possible that reduction of inflammation via corticosteroid treatment could have interfered with the remyelination process by slowing clearance of myelin debris, which is a prerequisite for OPC recruitment (36) and differentiation (37, 38), thereby negating any potential direct benefit of corticosteroids on remyelination.

Sex effect on remyelination was also investigated, with results suggesting that males had more remyelinating lesions than females. However, this result must be viewed with caution, since the two female animals (M#5 and M#6), twin pair, had shorter and more aggressive disease course compared to the other animals studied here. Furthermore, the lesions on these two animals were mainly ≤6 weeks old, possibly too little for substantial repair to have occurred.

The main limitation of the study is the small sample size. Although a higher number of animals would strengthen the study, as nonhuman primates are a scarce resource, we were able to maximize their use by focusing on individual lesions, rather than on number of animals developed remyelination. Another limitation is that different EAE immunization protocols were applied among marmosets. However, we did not observe any difference in lesion outcome across the different immunization schemes. Notably, previous studies from our group with a similar variety of EAE induction methods have not shown notable differences in disease course or lesion pathobiology (14, 24).

In conclusion, our study clearly demonstrates *in vivo* remyelination in the marmoset EAE model, further highlighting the utility of this animal model in investigating the repair of inflammatory demyelination. Remyelination was reliably detected and discriminated using serial *in vivo* MRI with automated segmentation. Our data indicate that the current model is an excellent avenue for investigating mechanisms of remyelination, and monitoring remyelination in a manner that is easily translatable to clinical studies in human beings.

## ACKNOWLEDGMENT AND COMPETING INTERESTS

This study was funded by the National Institute of Neurological Disorders and Stroke Intramural Research Program and Vertex Pharmaceuticals. We thank Dr. Tianxia Wu for statistical analyses. We thank everyone who took part in taking care of the animals and helped with acquisition of MRI scans.

---

**Supplementary Figure 1:**
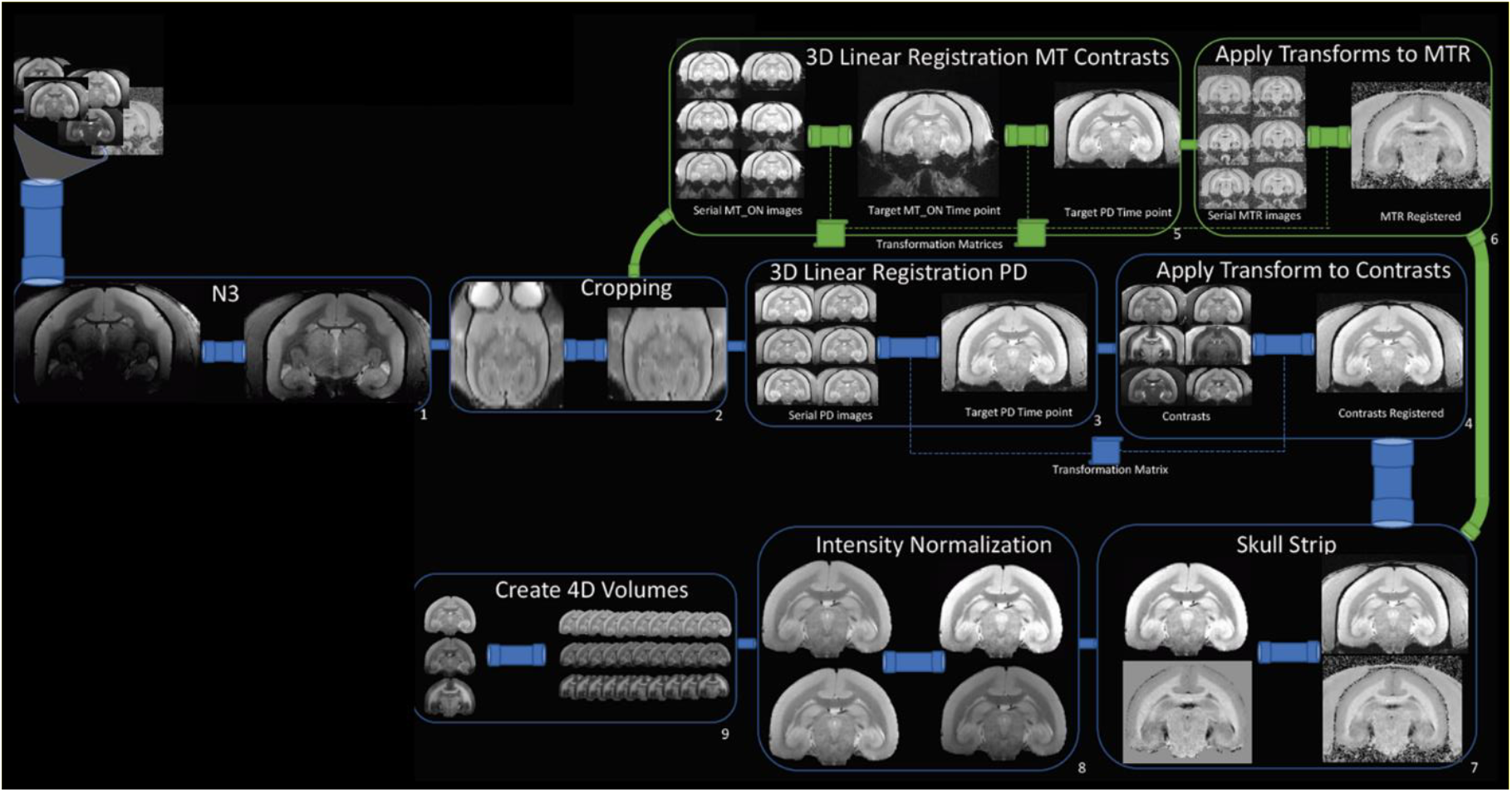
Workflow of the MRI post-processing pipeline. PD = proton density; MT = magnetization transfer; MTR = magnetization transfer ratio; N3 = bias field correction.

**Supplementary Figure 2:**
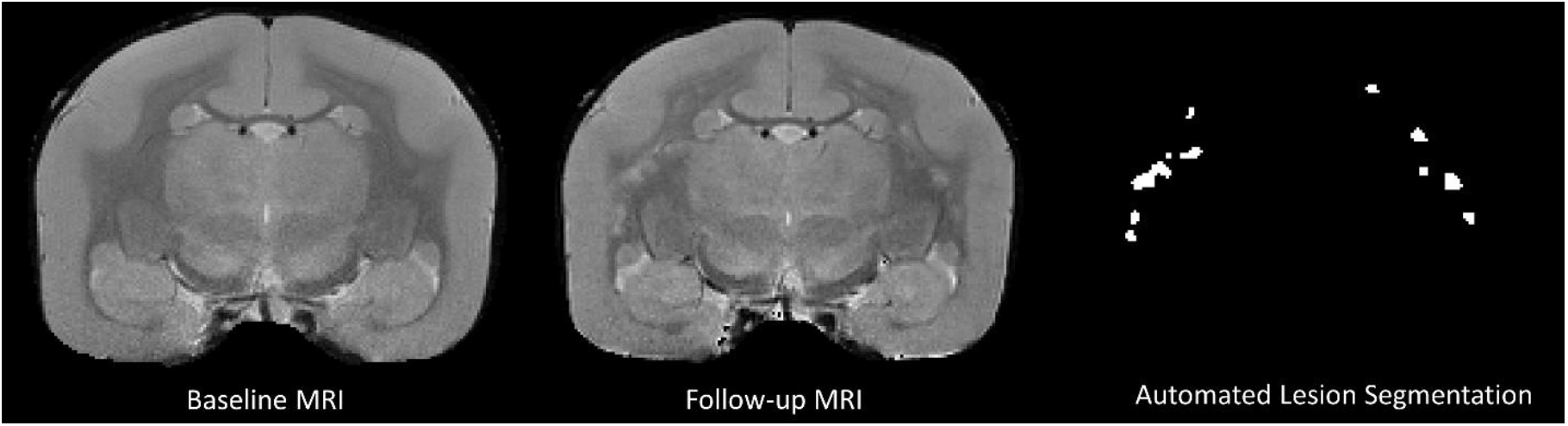
Post-processed proton-density weighted images used as input for the convolutional neural network -based lesion segmentation. All hyperintense lesions that appeared on the follow-up MRI performed post-immunization (M#1, 22 weeks post-immunization) were captured by the automated algorithm.

**Supplementary Table 1.** Immunohistochemistry methodology. For each of the immunohistochemical targets, respective companies, clonalities and hosts, and methods for antigen retrieval, blocking, and primary and secondary antibody inoculation are listed. HIER = heat-induced epitope retrieval; RT = room temperature; P = polyclonal antibody; M = monoclonal antibody; Rb = rabbit; Ms = mouse

|  | Company | Host / Clon-<br>ality | Antigen Retrieval<br>Method | Protein Blocking Rea-<br>gent | Primary Anti-<br>body Concen-<br>tration | Primary Anti-<br>body Inocula-<br>tion | Secondary Anti-<br>body |
| --- | --- | --- | --- | --- | --- | --- | --- |
| *Iba1* | Wako | Rb, P | HIER with Citrate<br>Buffer, 20" | Protein Block (Dako),<br>20" for HRP; 2.5% Horse<br>Serum (Vector) for AP | 1:400 | 1 hour, RT | Powervision<br>Poly-HRP or<br>ImmPRESS AP |
| *PLP* | BioRad | Ms, M | HIER with Citrate<br>Buffer, 20" | Protein Block (Dako),<br>20" | 1:200 | 1 hour, RT | Powervision<br>Poly-HRP |
| *ASPA* | GeneTex | Rb, P | HIER with Citrate<br>Buffer, 45" | Protein Block (Dako),<br>20" | 1:1500 | 1 hour, RT | Powervision<br>Poly-HRP |
| *Olig2* | EMD<br>Millipore | Rb, P | None when dou-<br>ble-staining | 2.5% Horse Serum (Vec-<br>tor), 20" | 1:200 | 1 hour, RT | ImmPRESS AP |

